# Purple bacteria as added-value protein ingredient in shrimp feed: *Litopenaeus vannamei* growth performance, and tolerance against *Vibrio* and ammonia stress

**DOI:** 10.1101/2020.02.25.964007

**Authors:** Abbas Alloul, Mathieu Wille, Piergiorgio Lucenti, Peter Bossier, Gilbert Van Stappen, Siegfried E. Vlaeminck

**Affiliations:** Research Group of Sustainable Energy, Air and Water Technology, Department of Bioscience Engineering, University of Antwerp, Groenenborgerlaan 171, 2020 Antwerpen, Belgium; Laboratory of Aquaculture & Artemia Reference Center, Faculty of Bioscience Engineering, Ghent University, Coupure Links 653, 9000 Gent, Belgium

**Keywords:** single-cell protein, microbial protein, purple phototrophic bacteria, acute hepatopancreatic necrosis disease, early mortality syndrome, fishmeal replacement

## Abstract

Purple non-sulfur bacteria (PNSB) biomass is an emerging alternative protein source, yet research of PNSB as added-value protein ingredient is limited. This research aimed to study the use of PNSB as protein source for shrimp and investigate the shrimp’s tolerance against *Vibrio* and ammonia stress. A 28-day shrimp feeding trial was performed with *Rhodopseudomonas palustris, Rhodobacter capsulatus* and a mixed PNSB culture. PNSB contained in feed (5-10% protein substitution) resulted in 5-26% higher individual weights, better feed conversions ratios (1.2-1.7) compared to commercial feed (1.7) and tolerance against ammonia. In parallel, the effect of PNSB on the growth of *Vibrio* pathogens was tested *in vitro.* The species *Rps. palustris, Rb. capsulatus, Rb. sphaeroides, Rhodospirillum rubrum* and *Afifella marina* suppressed the growth of *Vibrio parahaemolyticus* TW01 and *V. campbellii* LMG 21363. Overall, this study demonstrated the potential of PNSB as nutritious feed ingredient for shrimp. This can contribute to circular economy, as PNSB enable resource recovery from wastewater.

Graphical abstract

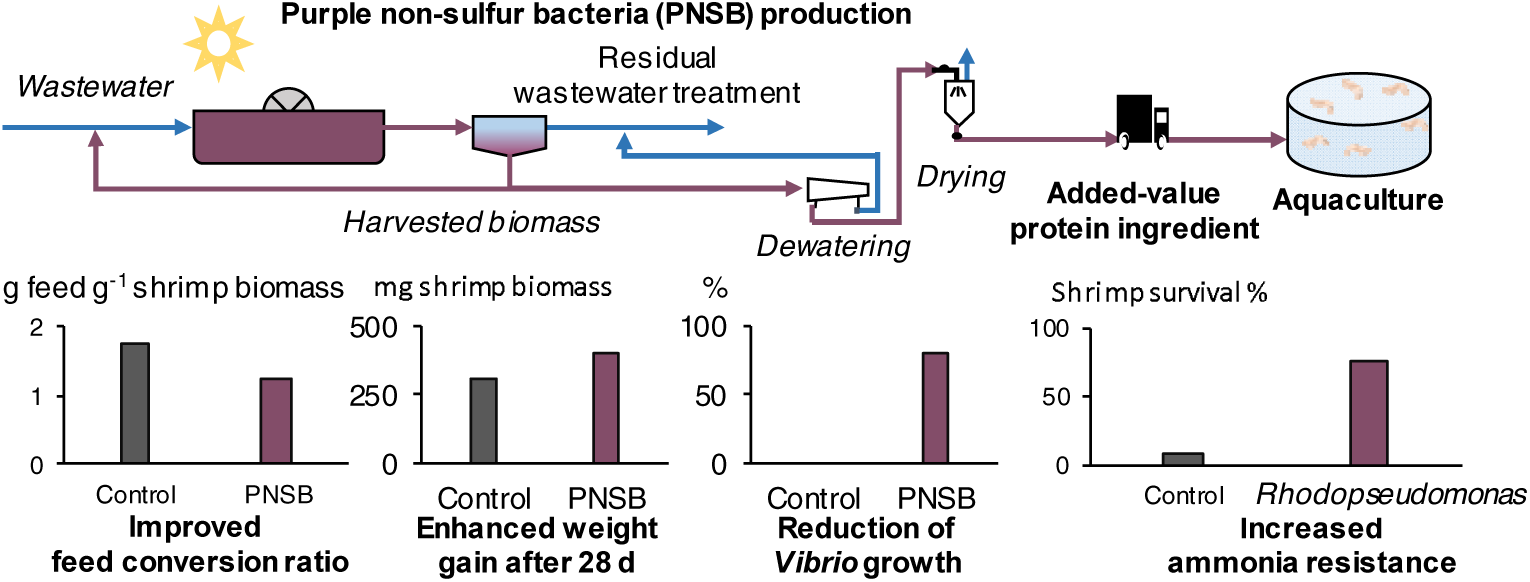

**Highlights:**

i. Purple non-sulfur bacteria (PNSB) enhance the growth performance of shrimp
ii. PNSB-fed shrimp has better feed conversion ratio, growth rate and weight gain
iii. Shrimp fed with *Rhodopseudomonas* are more resistance against ammonia stress
iv. Freeze-dried and live PNSB inhibit *Vibrio* pathogens *in vitro*

## 1 Introduction

Aquaculture is the fastest growing food production sector, expected to provide 60% of the fish available for human consumption by 2030 (FAO, 2016). The most-traded aquaculture products for decades were shrimp, comprising 15% of the total production (FAO, 2018). Shrimp cultivation and aquaculture, in general, depend on protein sources for aquafeeds, such as fishmeal and soybean meal, which have a high environmental impact (FAO, 2018; Fearnside, 2001). Finding alternative protein sources is, therefore, a prerequisite to decrease the dependency on conventional protein and guarantee a more sustainable growth of the aquaculture industry (Frost & Sullivan, 2018).

The use of microbial biomass as a dietary protein source for feed, so called microbial protein or single-cell protein, offers a sustainable alternative (Spiller et al., 2019). Microbial protein produced on wastewater will further increase sustainability by eliminating water and nutrient (C, N and P) requirements. However, production costs for microbial protein, even produced on wastewater (Alloul et al., 2020; Alloul et al., 2019), will still be higher than the market price of fishmeal (€ 2 kg^-1^ protein) or soybean meal (€ 0.7 kg^-1^ protein; IndexMundi, 2019). Therefore, added-value properties beyond purely the nutritional content of microbial protein, such as growth and health improvement for shrimp, ought to be investigated to increase the value of the product.

Purple non-sulfur bacteria (PNSB) are promising microorganisms that can be used as source of microbial protein (Alloul et al., 2018; Alloul et al., 2019). There biomass shows high potential as nutritious feed as it is rich in essential vitamins and carotenoid pigments (Sasaki et al., 1998). These phototrophic bacteria can be produced highly selectively on wastewater for proteins due to their ability to grow under anaerobic conditions in the light (Hülsen et al., 2016a; Hülsen et al., 2016b; Puyol et al., 2017). To date, studies of PNSB as protein substitution in shrimp feed are limited. Moreover, it is not known if PNSB contained in feed can be of added-value as growth promotor or enhance the tolerance of shrimp against *Vibrio* pathogens or ammonia stress. One of the few studies that has explored PNSB as shrimp feed, investigated their effect at low protein substitution levels (1.2-5.8%) and mainly looked to immune parameters (Chumpol et al., 2018). Hence, it could be argued that this research was more a probiotic than a protein replacement study. Studies have also shown that PNSB have anti-*Vibrio* properties when administered live (not as feed) to shrimp ponds (Chumpol et al., 2017; Wang & Gu, 2010). In addition, *in vitro* agar plate experiments revealed that some PNSB strains of *Rhodobacter sphaeroides* and *Afifella marina* have anti-*Vibrio* properties (Chumpol et al., 2017). However, thus far, no studies are available that investigate the anti-*Vibrio* potential of PNSB contained in feed.

This study has the objective to investigate PNSB as an added-value protein ingredient for shrimp. Therefore, a 28-day postlarval shrimp feeding trial was performed with *Rhodopseudomonas palustris* and *Rhodobacter capsulatus* at two inclusion levels along with a mixed culture holding *Rb. capsulatus* as a proxy for PNSB produced on wastewater. The feeding trial was followed by an ammonia stress test to validate if dietary inclusion of PNSB could improve fitness/stress and a *Vibrio* challenge to validate the pathogenic resistance of shrimp fed with PNSB. An *in vitro* screening of several PNSB species was performed in parallel to understand the effect of type of metabolism (i.e. photoheterotrophic, chemoheterotrophic and photoautotrophic) and live or freeze-dried PNSB administration on the growth of two *Vibrio* pathogens.

## 2 Materials and methods

### 2.1 Shrimp feeding trial

#### 2.1.1 Cultivation and preparation of PNSB

*Rps. palustris* LMG 18881 and *Rb. capsulatus* isolated by Alloul et al. (2019) were phototrophically cultivated in 500 mL flasks. A volatile fatty acid medium adapted from Alloul et al. (2019) was used, containing a 1/1/1 mixture of acetate, propionate and butyrate at a chemical oxygen demand (COD) concentration of 3 g L^-1^. Illumination was done with two halogen lamps (Sylvania, Germany) at a light intensity of 36 W m^-2^. The flasks were stirred with a multipoint stirrer at 300 rpm (Thermo Scientific, USA) and the temperature was 28°C. The broth was collected in 20 L tanks after 72 h of cultivation, leaving 50 mL of inoculum for a new batch. The collected broth was then centrifuged (Beckman Coulter, USA) at 7,300 g for 5 min. Subsequently, pellets were washed two times with distilled water to remove salts and centrifuged again. The washed pellets were collected in 50 mL falcon tubes and freeze-dried (Thermo Scientific, USA) for 48 h. After drying, PNSB powders were ground with a mortar and stored at −20°C to be used for the diet formulation.

A raceway reactor (MicroBio Engineering, USA) was also operated to produce a mixed PNSB culture holding *Rb. capsulatus* at an abundance between 14-56% as a proxy for PNSB produced on wastewater (Alloul et al., 2020). The 100 L reactor with a surface volume ratio of 5 m^3^ m^-2^ was operated at a sludge retention time of 2 d, using the same volatile fatty acid medium as used for the pure cultures. Temperature and pH were respectively controlled at 28°C and 7. A halogen lamp (Sylvania, Germany) was used to illuminate the reactor at a light intensity of 54 W m^-2^ and an illumination period of 12h-light/12h-dark. The broth was stirred through a paddlewheel at a speed of 30 rpm. The produced PNSB were then centrifuged, washed and freeze-dried using the same procedure as described for the pure cultures. Table 1 presents an overview of the proximate analyses for all cultivated PNSB.

**Table 1.** Proximate analysis with averages ± standard deviation (n=3) for two pure cultures and a mixed culture holding *Rb. capsulatus*. Dry weight (DW), Markwell protein, crude protein, lipids and ash analyzed in the laboratory of the Research Group of Sustainable Energy, Air and Water Technology.

| **PNSB species** | *Rps. palustris* | *Rb. capsulatus* | *Rb. capsulatus* |
| --- | --- | --- | --- |
| **Culture** | Pure | Pure | Mixed |
| Dry weight (g 100 g<sup>-1</sup> product) | 79.4 ± 1.1 | 81.6 ± 0.4 | 89.7 ± 1.7 |
| Markwell protein (g 100 g<sup>-1</sup> DW) | 48.1 ± 3.5 | 48.0 ± 3.1 | 60.5 ± 0.2 |
| Crude protein (g 100 g<sup>-1</sup> DW) | 67.0 ± 1.6 | 66.9 ± 5.9 | 73.7 ± 5.5 |
| Lipid (g 100 g<sup>-1</sup> DW) | 14.7 ± 5.3 | 14.6 ± 7.6 | 13.1 ± 2.2 |
| Ash (g 100 g<sup>-1</sup> DW) | 35.8 ± 2.4 | 36.4 ± 3.3 | 17.1 ± 2.8 |

#### 2.1.2 Diet preparation

The experiment consisted of six practical diets, based on a commercial formulation, with different substitution levels of total protein with PNSB: (i) a commercial formulation without PNSB close to what would be used in the field as experimental control, (ii & iii) *Rps. palustris* at a protein substitution level of 5% and 11%, (iv & v) *Rb. capsulatus* at a protein substitution level of 5% and 11% and (vi) a mixed culture containing *Rb. capsulatus* at a protein substitution level of 11%. All diets were formulated to be equivalent for levels of protein (35%) and fat (7%). Formulation of the diets was done by VDS (Deerlijk, Belgium) with the Mevoblend software designed by AMS Software (Roeselare, Belgium). All ingredients used were industrial grade (VDS, Deerlijk, Belgium), except for the binder carboxymethyl cellulose (VWR Chemicals, Belgium, Oud-Heverlee, product 22525296). The diets were prepared by mixing all the ingredients in a kitchen machine (Kenwood Major Titanium KMM060) and adding water to obtain a firm dough. The dough was then passed through a meat grinder (Kenwood Major Titanium KMM060) with a hole size of 4 mm. The diet strands were subsequently dried for 48 h at room temperature. After drying, the diets were ground (Kenwood Major Titanium KMM060) and sieved to make four pellet fractions: (i) 300-500 µm, (ii) 500-800 µm, (iii) 800-1000 µm and (iv) 1000-1200 µm. Pellets were then packed and stored at 4°C. The ingredient composition and proximate analyses are presented in Table 2.

**Table 2.** Ingredient composition and proximate analysis of diets for *Litopenaeus vannamei*. All ingredients were industrial grade from VDS (Deerlijk, Belgium) except for carboxymethyl cellulose (CMC) and purple non-sulfur bacteria (PNSB). Dry weight (n=3), Markwell protein (n=3), lipids (n=4) and ash (n=3) analyzed in the laboratory of the Research Group of Sustainable Energy, Air and Water Technology. Crude protein (n=2) analyzed in the laboratory of Aquaculture & Artemia Reference Center.

| PNSB species | Control<br>feed | *Rps.*<br>*palustris* | *Rps.*<br>*palustris* | *Rb.*<br>*capsulatus* | *Rb.*<br>*capsulatus* | *Rb.*<br>*capsulatus* |
| --- | --- | --- | --- | --- | --- | --- |
| Culture | - | Pure | Pure | Pure | Pure | Mixed |
| Protein substitution | 0% | 5% | 11% | 5% | 11% | 11% |
| Ingredients (g 100 g<sup>-1</sup> product) |  |  |  |  |  |  |
| PNSB | 0 | 5 | 10 | 4.6 | 9.5 | 7.0 |
| Wheat meal | 31.0 | 30.2 | 38.4 | 30.4 | 39.0 | 30.1 |
| Soybean meal | 38.0 | 38.0 | 17.2 | 38.0 | 17.1 | 37.4 |
| Fishmeal | 20.0 | 17.4 | 27.7 | 17.3 | 27.6 | 15.0 |
| Methionine | 0.0 | 0.05 | 0.0 | 0.05 | 0.0 | 0.0 |
| Tuna oil | 2.5 | 2.2 | 0.8 | 2.2 | 0.9 | 2.2 |
| Crustacean premix | 0.5 | 0.5 | 0.5 | 0.5 | 0.5 | 0.5 |
| Ca(H<sub>2</sub>PO<sub>4</sub>)<sub>2</sub> | 1.2 | 1.2 | 1.2 | 1.2 | 1.2 | 1.2 |
| CaCO<sub>3</sub> | 1.5 | 1.5 | 0.2 | 1.5 | 0.2 | 1.5 |
| CMC** | 2.5 | 2.5 | 2.5 | 2.5 | 2.5 | 2.5 |
| Lecithin | 1.5 | 1.5 | 1.5 | 1.5 | 1.5 | 1.5 |
| Proximate analysis |  |  |  |  |  |  |
| Dry weight<br>(g 100 g<sup>-1</sup> product) | 93.24 ± 0.51 | 92.65 ± 0.37 | 90.48 ± 1.26 | 93.68 ± 0.14 | 92.38 ± 0.2 | 93.77 ± 1.02 |
| Markwell protein<br>(g 100 g<sup>-1</sup> DW) | 36.22 ± 0.2 | 36.15 ± 3.95 | 35.72 ± 0.44 | 36.37 ± 2.02 | 38.03 ± 0.5 | 36.94 ± 1.12 |
| Crude protein<br>(g 100 g<sup>-1</sup> DW) | 37.86 ± 0.55 | 39.21 ± 0.28 | 39.75 ± 0.22 | 38.32 ± 0.37 | 46.76 ± 0.19 | 37.79 ± 0 |
| Lipids<br>(g 100 g<sup>-1</sup> DW) | 7.52 ± 0.47 | 7.79 ± 0.78 | 7.09 ± 1.28 | 7.34 ± 0.47 | 9.06 ± 0.31 | 7.48 ± 0.84 |
| Ash<br>(g 100 g<sup>-1</sup> DW) | 12.87 ± 0.68 | 11.55 ± 0.65 | 11.13 ± 3.58 | 10.9 ± 0.27 | 13.54 ± 0.29 | 10.79 ± 0.79 |

#### 2.1.3 Shrimp source and experimental design

*Litopenaeus vannamei* were imported as postlarvae from Shrimp Improvement Systems (Florida, USA). These shrimp are certified to be specific pathogen free for the following pathogens: WSSV, YHV/GAV/LOV, TSV, IHHNV, BP, MBV, BMN, IMN, Microsporidians, Haplosporidians and NHP bacteria. Upon arrival, shrimp postlarvae were transported to the facilities of ARC and reared into a recirculating system containing natural seawater at a salinity of 35 g L^-1^ and a pH of 7.8-8.1. They were initially fed Artemia (live feed) and after that weaned onto a commercial pelleted feed (Crevetec BVBA, Belgium).

The feeding trial was conducted for 28 days in 24 square grey PVC tanks with a useful volume of 30 L and a surface area of 0.135 m^2^. Temperature and illumination were controlled at respectively 28-29°C and 12h-light/12h-dark. All 24 tanks had a stocking density of 100 shrimps per tank. Experimental tanks were arranged in a randomized block design. Four independent recirculation units were set up. Each unit contained six tanks connected to a submerged biofilter tank and a sump tank from where the water was pumped back into an overhead tube, supplying the shrimp tanks with filtered water. From the shrimp tanks, water then flowed back to the biofilter by gravity. The water recirculation rate was 700% of the tank volume per day. Approximately 10% of fresh seawater was added to compensate for evaporation and water loss due to siphoning.

Every dietary treatment was replicated four times, in which one replicate of every treatment was distributed randomly among one of the four recirculation systems (randomized block). Shrimp were fed three times per day at 9h, 13h and 17h. The feed ration was calculated based on a feed conversion ratio (FCR) of 1.6 g feed g^-1^ shrimp biomass and a specific growth rate (SGR) of 15% d^-1^. A sample of roughly 20 shrimp per tank was weighed weekly to check actual SGR and to adjust the feed dosage when required. The remaining feed and feces were siphoned out daily to maintain water quality. After four weeks, all shrimps were weighed and counted to determine the growth parameters and survival. Growth parameters were calculated as followed: weight gain = (final weight − initial weight), SGR = (100 × (ln final weight − ln initial weight)/trial duration), FCR = (feed intake/weight gain), survival = (final number shrimp/initial number of shrimp) × 100.

All water parameters were measured daily. Salinity was measured with a handheld refractometer (model, 635-0636, VWR International BVBA, Belgium) and temperature with a liquid in glass thermometer. NH_4_ ^+^-N and NO_2_ ^-^-N were measured through standard kits (Tetra, Germany).

### 2.2 Exploring the anti-*Vibrio* potential of PNSB

#### 2.2.1 PNSB and *Vibrio* species

In parallel with the shrimp feeding trial, *in vitro* experiments were performed to explore the anti-*Vibrio* potential of PNSB. In addition, a post-feeding stress and challenge tests were performed to validate if dietary inclusion of PNSB could improve fitness/stress resistance and to explore the resistance of shrimp fed with PNSB against *Vibrio* pathogens. These *Vibrio* pathogens are the causing agents for acute hepatopancreatic necrosis disease (AHPND), one of the most devastating diseases in the shrimp sector (Tran et al., 2013). It affects shrimp postlarvae within 20-30 days causing up to 100% mortality and results in enormous economic repercussions.

Five PNSB species were used throughout these experiments: (i) *Rhodopseudomonas palustris* LMG 18881, (ii) *Rhodobacter sphaeroides* LMG 2827, (iii) *Rhodospirillum rubrum* S 1H, (iv) *Afifella marina* BN 126 and (v) *Rb. capsulatus* isolated by Alloul et al. (2019). Two virulent pathogens were selected for the study : (i) *Vibrio parahaemolyticus* TW01 obtained from AHPND diseased shrimp in Thailand and (ii) *V. campbellii* LMG 21363 (Vanmaele et al., 2015).

#### 2.2.2 96-Well plate experiments with freeze-dried PNSB

96-Well plate experiments were performed to investigate the effect of freeze-dried PNSB on the growth of *Vibrio* pathogens. PNSB deployed for wastewater treatment will always be cultivated photoheterotrophically. However, PNSB are also able to use other metabolisms. Therefore, all five PNSB species described above were explored on two or three of their metabolisms: (i) photoheterotrophic, (ii) photoautotrophic and (iii) chemoheterotrophic. *Dunaliela salina* SAG 184.80 was also tested as its antimicrobial properties against *Vibrio* pathogens have already been reported (Lustigman, 1988). A consortium of aerobic heterotrophic bacteria grown on synthetic brewery wastewater (Supplementary Material S1) was also included in the experiment.

PNSB cultivated photoheterotrophically were grown anaerobically in the light as described in section 2.1.1 on the medium based on Alloul et al. (2019). For their photoautotrophic cultivation, the carbon source in this medium was replaced by hydrogen gas (electron donor) in the headspace and the flasks were incubated in the light at 28°C. To grow PNSB chemoheterotrophically, the volatile fatty acid mixture was replaced by fructose at a COD concentration of 3 g L^-1^ and the bacteria were incubated aerobically in the dark at 28°C. *D. salina* was cultivated using a modified Johnson’s medium (Sui & Vlaeminck, 2019). All types of biomass were eventually centrifuged at 7,300 g for 5 min and washed two times with distilled water. The cells were then dried in a freeze dryer (Thermo Scientific, USA) for 48 h. After drying, 2 mL of absolute ethanol was added to the pellets to kill-off and prevent the growth of PNSB. The pellets were then placed under the laminar flow in order to evaporate the ethanol.

The 96-Well plate experiments were performed in a working volume of 150 µL with Marine Broth (Carl Roth, Germany). *V. parahaemolyticus* TW01 and *V. campbellii* LMG 21363 were added to the wells at a concentration of 10^5^ cells mL^-1^ (OD_550nm_ of 1.0 corresponds to 1.2 × 10^9^ cells mL^-1^; Niu et al., 2014). The PNSB, *D. salina* and the aerobic heterotrophic bacteria were then dosed to the wells at a volatile suspended solids concentration of 0.5 g L^-1^. Several controls were included in the test: (i) *Vibrio* pathogens without PNSB to test their natural growth in the wells. This growth rate was used to determine the growth rate reduction (Figure 1); (ii) PNSB without pathogens to check whether the freeze-dried and ethanol treated biomass was still revivable.; (iii) Marine Broth without pathogens and without PNSB was also incubated to test for contamination of the medium.; (iv) *Vibrio* pathogens were grown with rifampicin at a concentration of 20 mg L-1 as a negative control. All tests were performed in triplicate. After adding of pathogens and PNSB, 96-Well plates were incubated in a microplate reader (Biotek, USA) at 28°C with vigorous shaking (282 rpm) for aeration. The growth was continuously monitored by measuring the absorbance at 660 nm. The *Vibrio* growth reduction was eventually calculated as follows: (Growth rate_*Vibrio*_*-*Growth rate_*Vibrio+*PNSB_)/Growth rate_*Vibrio*_ x 100.

**Figure 1.**
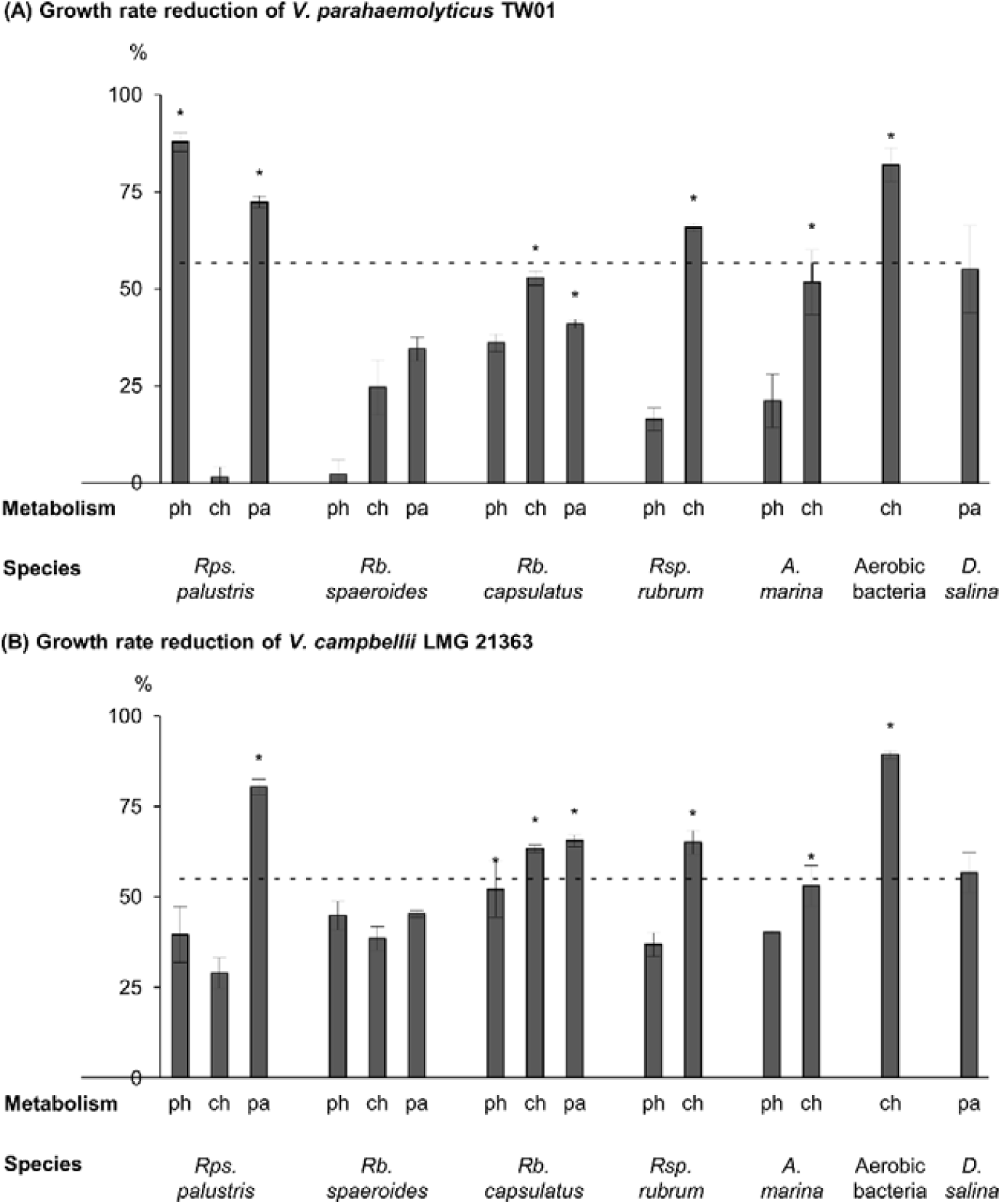
Percentage of growth rate reduction for (A) *Vibrio parahaemolyticus* TW01 and (B) *V. campbellii* LMG 21363 by supplementation of purple non-sulfur bacteria, a consortium of aerobic bacteria or a microalga. Microorganisms were grown photoheterotrophically (ph), chemoheterotrophically (ch) and photoautotrophically (pa). Error bars show standard deviation (n=3). The dotted line shows the average of *Dunalialla salina* because the results were statistically compared with this microalga. * Treatments that were significantly higher (*P* < 0.05) or equal to (*P* > 0.05) *Dunaliella salina*.

#### 2.2.3 Agar spot assay with live PNSB

Experiments in section 2.2.2 were performed to unravel the effect of freeze-dried PNSB on *Vibrio* growth. This section had the objective to study the effect of live administered PNSB on *Vibrio* growth.

The media described in 2.2.2 (i.e. photoheterotrophic, photoautotrophic and chemoheterotrophic) were supplemented with agar at 15 g L^-1^ and poured in square agar plates. The five PNSB species described in 2.2 were first grown photoheterotrophically and 2µL of the inoculum was then spotted in triplicate on the three types of agar plates. As a negative control, 2 µL of filtered supernatant (0.22 µm) was spotted on the agar plates to verify whether it affected the pathogens. Subsequently, agar plates were incubated at 28°C according to their respective metabolism: (i) photoheterotrophically in an anaerobic bag flushed with nitrogen gas illuminated with a halogen lamp (Sylvania, Germany, (ii) photoautotrophically in an anaerobic bag flushed with hydrogen gas (electron donor) illuminated with a halogen lamp (Sylvania, Germany) and (iii) chemoheterotrophically in the dark. After 48 h of incubation, agar plates were overlaid with soft agar (5 g L^-1^) containing either *V. parahaemolyticus* TW01 or *V. campbellii* LMG 21363 at a concentration of 10^5^ cells mL^-1^. After 48 h of incubation, zones of inhibition were measured from the edge of the PNSB colony until the end of the zone. The supernatant of all five PNSB broths was also tested as a negative control.

#### 2.2.4 Ammonia stress test and *Vibrio* challenge

After the feeding trial, ammonia stress and *Vibrio* challenge test was performed according to the experimental design of Le Hong (2008). These tests were performed to validate if dietary inclusion of PNSB could improve fitness/stress resistance and explore the resistance of shrimp fed with PNSB against *Vibrio* pathogens.

Shrimp from four replicates of each diet were pooled and randomly distributed into disposable plastic boxes with a volume of 1 L, filled with 500 mL of fresh seawater. Each box was stocked with one shrimp to exclude cannibalism. Boxes were covered to prevent cross contamination and shrimp from jumping out. Shrimp of every diet were exposed to ammonia stress, a *V. parahaemolyticus* TW01 challenge test and a *V. campbellii* LMG 21363 challenge test. These *Vibrio* pathogens were used because they are recognized to be virulent (Vanmaele et al., 2015). All challenge tests were performed in duplicate (ammonia stress) or triplicate (*Vibrio* challenge tests). Each replicate consisted out of eight boxes with each box containing one shrimp. Thus for a triplicate, a total of 24 shrimp were used and for a duplicate 16 shrimp. All tests were performed in a temperature (28 °C) controlled room. As a negative control, shrimp fed with the control feed were not exposed to the pathogens. Feeding of shrimp was continued throughout the challenge test with their specific diet by adding one grain (800-1000 µm) to the boxes.

For the ammonia stress challenge, a range-finding test was first performed by exposing shrimp fed with the control feed (i.e. 0% PNSB substitution) to NH_3_-N concentrations of 0, 0.5, 1, 1.7, 3 and 5.3 mg N L^-1^ (addition of NH_4_Cl at pH of 8). The survival was recorded after 24 h and 48 h. From these data, a 48h-LC_50_ value was derived using the program Excel Microsoft Office Professional Plus 2016 (Supplementary Material S2). This LC_50_, which corresponded to 3 mg N L^-1^, was then used to evaluate the fitness of animals from the different groups in the actual ammonia stress tests. Survival was monitored after 24 h and 48 h. As a negative control, shrimp fed with the control feed were not exposed to ammonia

For the pathogen challenge tests, *V. parahaemolyticus* TW01 and *V. campbellii* LMG 21363 were initially grown on Marine Agar at 28°C for 24h. A colony was then transferred to Marine Broth and incubated at 28 °C for 24h with continuous shaking. Prior to the experiment, pathogens were again diluted 10 times with Marine Broth at 28°C to guarantee that the *Vibrio* were in exponential phase. The *V. parahaemolyticus* TW01 and *V. campbellii* LMG 21363 were then transferred to plastic boxes with the individual shrimp at a concentration of respectively 10^6^ cells mL^-1^ and 10^7^ cells mL^-1^. Survival was monitored every 12-14 h for a period of maximal 5 days. The shrimp tested with *V. campbellii* LMG 21363 were simultaneously exposed to an ammonia concentration of 35 mg N L^-1^ (corresponding to the 24h-LC_10_ of the control animals) to make the shrimp more susceptible to the pathogen. After 62 h, shrimp of the *V. campbellii* LMG 21363 challenge were almost not affected. Therefore, an extra dosage of 10^7^ cells mL^-1^ was added to the boxes. Mortality rates (%mortality h^-1^) were determined by calculating the slope between mortality and time.

### 2.3 Analytic procedures

The dry weight and water content of the PNSB biomass and the six practical diets were determined gravimetrically in triplicate or quadruplicate by drying at 105°C for 24h. After drying, samples were incinerated at 550°C for 2 h to determine the ash content. Total lipid content of the different PNSB cultures and the six diets were also determined gravimetrically according to Bligh and Dyer (1959) in triplicate or quadruplicate. Protein content was analyzed by Markwell et al. (1978) in triplicate, an adapted Lowry et al. (1951) protocol and through a Kjeldahl nitrogen measurement in quadruplicate.

### 2.4 Statistical analyses

The analysis of variance test and post-hoc pairwise comparisons using the Tukey’s range test were conducted for multiple comparisons. Normality was tested using the Shapiro-Wilk normality test and homogeneity of variances using a Levene’s test. If normality was rejected, the non-parametric Kruskal-Wallis rank sum test and post-hoc pairwise comparisons using the Mann-Whitney U test (*p-*values were adjusted using the Benjamini-Hochberg correction) were performed. If homoscedasticity was rejected or in case of unequal sample size, the Welch’s t-test was conducted. Analysis of covariance was performed to compare regression lines. A significance level of *p* < 0.05 was chosen. All analyses were performed in R (version 3.4.1) using RStudio (RStudio®, USA) for Windows.(R Core Team, 2017).

## 3 Results and discussion

### 3.1 PNSB contained in feed enhances the growth performance of shrimp

Finding alternative protein sources for shrimp production is key for a more sustainable industry (Frost & Sullivan, 2018). Therefore, this feeding trial had the objective to demonstrate that PNSB are a suitable alternative for conventional protein ingredients.

The results of the feeding trial demonstrated that PNSB substitution resulted in similar and even enhanced growth performance of shrimp compared to a commercial feed (Table 3). SGR were on average 13.2 ± 2.7 % d^-1^, comparable to values in the field (Hung & Quy, 2013). Only *Rps. palustris* at a protein substitution of 11% resulted in a significantly higher SGR (13.5 ± 0.4 % d^-1^; *P*< 0.05) compared to the control feed (12.6 ± 0.5 % d^-1^). In terms of feed conversion ratio (FCR), *Rps. palustris* at a protein substitution of 5%, *Rb. capsulatus* at a protein substitution of 5% and 11% showed significantly better FCR (1.2-1.4 g feed g^-1^ shrimp biomass) compared to the control (1.7 ± 0.2 g feed g^-1^ shrimp biomass). Hence, shrimp fed with these experimental diets were more efficient in feed conversion compared to the control feed. All other diets had an equal FCR as the control feed (Table 3). Final individual weights showed again significantly higher values for *Rps. palustris* at a protein substitution of 5% and *Rb. capsulatus* at a protein substitution of 5% and 11% (Table 3). Survival after 28 days was overall 87 ± 4%, with similar performance for all experimental diets and the control feed (*p* > 0.05).

**Table 3.**
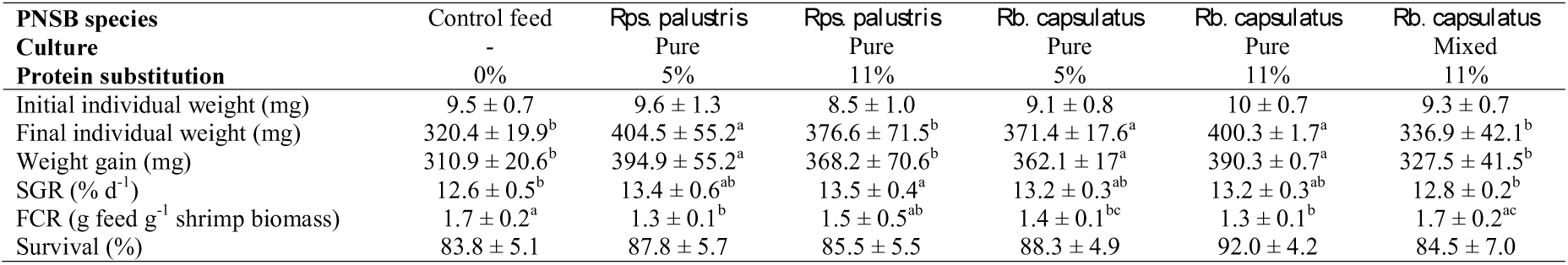
*Litopenaeus vannamei* feeding trial with a commercial feed (control) and purple non-sulfur bacteria (PNSB) at two protein substitution levels. Values show averages ± standard deviation for control feed (n=4), *Rhodopseudomonas palustris* 5% (n=4) and *Rhodopseudomonas palustris* 11% (n=4), *Rhodobacter capsulatus* 5% (n=3) and *Rhodobacter capsulatus* 11% (n=2) and *Rhodobacter capsulatus* 11% (mixed culture; n=4). Letters within columns indicate significant differences. A significance level of p < 0.05 was chosen.

| **PNSB species** | Control feed | *Rps. palustris* | *Rps. palustris* | *Rb. capsulatus* | *Rb. capsulatus* | *Rb. capsulatus* |
| --- | --- | --- | --- | --- | --- | --- |
| **Culture** | - | Pure | Pure | Pure | Pure | Mixed |
| **Protein substitution** | 0% | 5% | 11% | 5% | 11% | 11% |
| Initial individual weight (mg) | 9.5 ± 0.7 | 9.6 ± 1.3 | 8.5 ± 1.0 | 9.1 ± 0.8 | 10 ± 0.7 | 9.3 ± 0.7 |
| Final individual weight (mg) | 320.4 ± 19.9<sup>b</sup> | 404.5 ± 55.2<sup>a</sup> | 376.6 ± 71.5<sup>b</sup> | 371.4 ± 17.6<sup>a</sup> | 400.3 ± 1.7<sup>a</sup> | 336.9 ± 42.1<sup>b</sup> |
| Weight gain (mg) | 310.9 ± 20.6<sup>b</sup> | 394.9 ± 55.2<sup>a</sup> | 368.2 ± 70.6<sup>b</sup> | 362.1 ± 17<sup>a</sup> | 390.3 ± 0.7<sup>a</sup> | 327.5 ± 41.5<sup>b</sup> |
| SGR (% d<sup>-1</sup>) | 12.6 ± 0.5<sup>b</sup> | 13.4 ± 0.6<sup>ab</sup> | 13.5 ± 0.4<sup>a</sup> | 13.2 ± 0.3<sup>ab</sup> | 13.2 ± 0.3<sup>ab</sup> | 12.8 ± 0.2<sup>b</sup> |
| FCR (g feed g<sup>-1</sup> shrimp biomass) | 1.7 ± 0.2<sup>a</sup> | 1.3 ± 0.1<sup>b</sup> | 1.5 ± 0.5<sup>ab</sup> | 1.4 ± 0.1<sup>bc</sup> | 1.3 ± 0.1<sup>b</sup> | 1.7 ± 0.2<sup>ac</sup> |
| Survival (%) | 83.8 ± 5.1 | 87.8 ± 5.7 | 85.5 ± 5.5 | 88.3 ± 4.9 | 92.0 ± 4.2 | 84.5 ± 7.0 |

This study was the first to conduct a shrimp feeding trial with PNSB contained in the feed at relatively high protein substitution levels (5-11% protein). According to the authors’ knowledge, only Chumpol et al. (2018) have investigated PNSB as protein substitution for conventional shrimp feed. However, the protein substitution level was only between 1.2-5.8% and the biomass increase was merely 8-12 times the initial weight compared to 33 times in our study. In addition, the diets were not designed by substituting protein in the feed, yet by adding PNSB to a ground commercial feed. Consequently, it could be argued that the study of Chumpol et al. (2018) was more a probiotic rather than a protein substitution study.

For other aquaculture species, higher PNSB protein substitution levels were already tested. Banerjee et al. (2000) substituted 40% of the protein in a commercial diet of tilapia (*Oreochromis niloticus*) and observed a higher weight gain, survival and SGR compared to the control feed. An experiment with *Rhodopseudomonas gelatinosa* administered to a goldfish (*Carassius auratus*) diet at a protein substitution level of 31% also resulted in a higher SGR and a higher weight gain (Noparatnaraporn et al., 1987). All previous studies on PNSB as protein ingredient were performed with pure cultures. Only recently, Delamare-Deboutteville et al. (2019) have tested mixed PNSB cultures. They observed a similar performance compared to a commercial feed for the Asian sea bass (*Lates calcarifer*) up to a protein substitution level of 21%. At 30% protein substitution, SGR and FCR were worse compared to the commercial feed. This was also observed for shrimp fed with *Arthrospira* (*Spirulina platensis*) at a protein substitution of 81% (Macias-Sancho et al., 2014). The effect of high PNSB protein substitutions (>11% protein) should also be explored for shrimp and research on PNSB as aquafeed needs to focus more on mixed PNSB cultures instead of pure cultures.

To date, research on PNSB as protein substitution in shrimp feed and aquafeeds, in general, is limiting. It is, therefore, challenging to understand the underlying mechanism behind the enhanced performance of the shrimps in terms of FCR and weight gain compared to the commercial diet. A hypothesis might be that PNSB in the feed were able to colonize the gastrointestinal tract and assist the digestion. Chumpol et al. (2017) for example observed that shrimp administered with live PNSB are able to colonize the gastrointestinal tract and assist the digestion by excreting proteases, lipases and amylases. Our study was performed with freeze-dried PNSB. According to a growth experiment in Marine Broth (Supplementary Material S3), PNSB were able to be revived after freeze-drying. Therefore, PNSB contained in the feed may have assisted the digestion. PNSB are also known to accumulate poly-β-hydroxybutyrate (Sangkharak & Prasertsan, 2007) and it has been demonstrated that these compounds enhance the growth performance of shrimp (Gao et al., 2019). Next to their ability to produce enzymes and accumulate poly-β-hydroxybutyrate, PNSB also contain vitamins and pigments, which are antioxidants (Sasaki et al., 1998). All these compounds may have contributed to an enhanced performance of PNSB relative to the control feed.

### 3.2 Exploring the anti-*Vibrio* potential of PNSB

#### 3.2.1 Freeze-dried and live PNSB inhibit *Vibrio* growth

In parallel with the shrimp feeding trial, *in vitro* experiments were performed to explore the effect of PNSB species, their growth metabolism (i.e. photoheterotrophic, chemoheterotrophic and photoautotrophic) and live or freeze-dried PNSB administration on the growth of *Vibrio* pathogens.

Freeze-dried administration of PNSB showed promising results. *Rps. palustris* grown photoheterotrophically imposed the highest growth rate reduction on *V. parahaemolyticus* TW01 (88 ± 2% Figure 1). For *V. campbellii, Rps. palustris* grown photoautotrophically induced the highest growth rate reduction (80 ± 2%). The type of metabolism did not show a consistent growth rate reduction for both *Vibrio* species (*p* > 0.05).

Overall, the results showed that *Rps. palustris* (photoheterotrophic), *Rb. capsulatus* (both chemoheterotrophic and photoautotrophic), *Rsp. rubrum* (chemoheterotrophic) and *A. marina* (chemoheterotrophic) were effective against both *Vibrio* pathogens. This was evident by executing a Pearson correlation test between the results of *V. parahaemolyticus* TW01 and *V. campbellii* LMG 21363 (significant correlation *p* < 0.05). Therefore, high growth rate reductions for one pathogen imposed by a certain PNSB cultivated on a certain metabolism also occurred for the other pathogen.

A consortium of aerobic heterotrophic bacteria cultivated on synthetic brewery wastewater was also tested. These bacteria reduced the growth rate of both *V. parahaemolyticus* TW01 and *V. campbellli* by respectively 82 ± 4% and 89 ± 1%. The suppression might be due to traces of yeast (*Saccharomyces cerevisiae*) in the biomass, originating from the medium (0.5 g L^-1^ yeast), which has already been shown to affect *V. campbellii* (Soltanian et al., 2007).

*D. salina* was also tested as a benchmark because the antimicrobial potential of this microalga has already been reported (Lustigman, 1988). Our results confirm these findings with a growth rate reduction of 55 ± 11% for *V. parahaemolyticus* TW01 (Figure 1.A) and 57 ± 5% for *V. campbellli* (Figure 1.B).

Agar spot assays were performed after the 96-Well experiment (Figure 2). Only *Rb. capsulatus* and *Rps. palustris* inhibited *Vibrio* growth. *Rb. sphaeroides, Rsp. rubrum* and *A. marina* were all overgrown by the pathogens. As for the freeze-dried PNSB experiments (Figure 1), the type of metabolism did not have an effect on the size of the zone of inhibition. In both the experiments with freeze-dried or live PNSB, *V. campbellii* LMG 21363 was more susceptible to inhibition than *V. parahaemolyticus* TW01 (*p* < 0.05).

**Figure 2.**
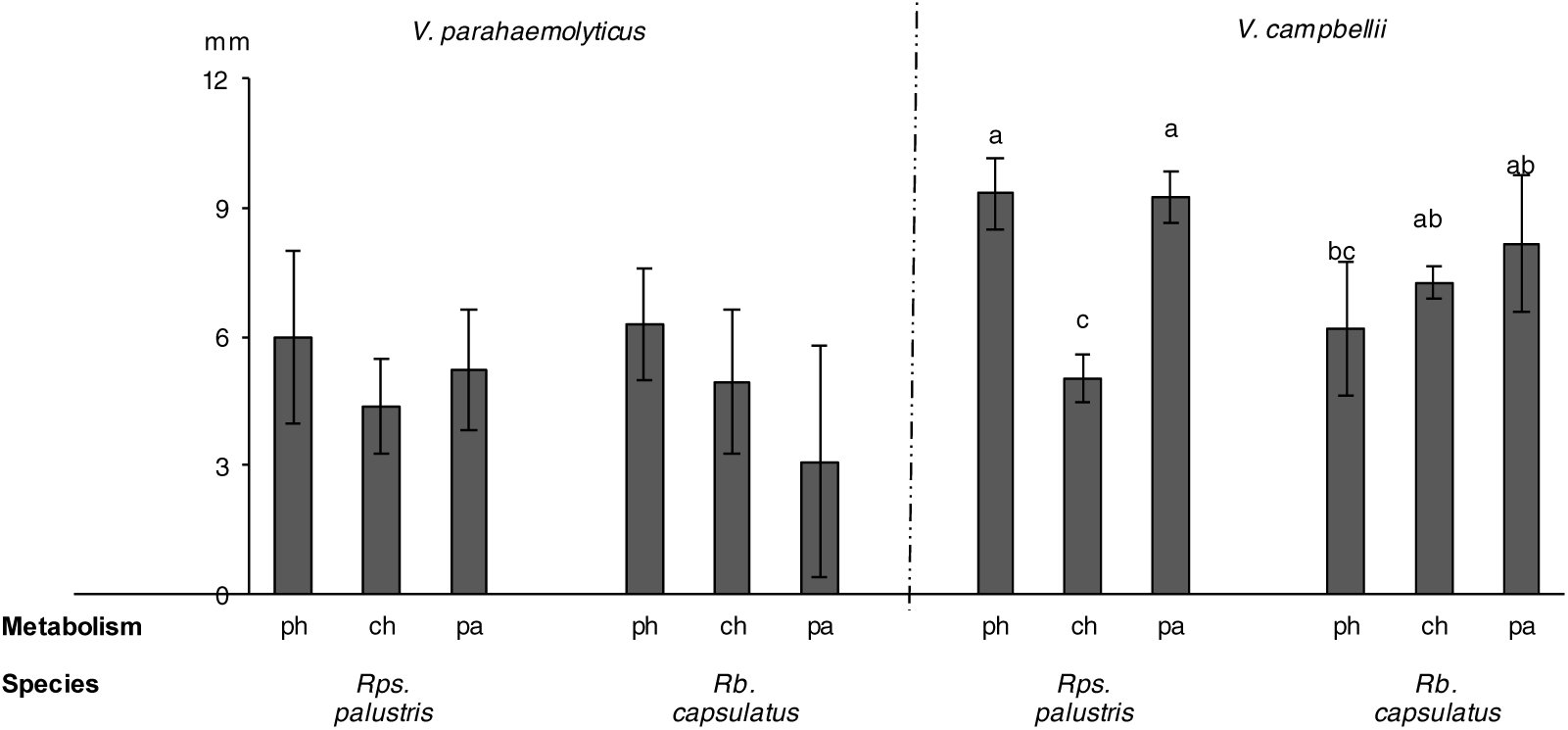
Zone of inhibition resulting from purple non-sulfur bacteria on the growth of *Vibrio parahaemolyticus* TW01 *and V. campbellii* LMG 21363. No zones of inhibition were observed for *Rhodobacter sphaeroides, Rhodospirillum rubrum* and *Afifella. marina.* Purple non-sulfur bacteria were grown photoheterotrophic (ph), chemoheterotrophic (ch) and photoautotrophic (pa). Error bars show standard deviation (SD; n=3). Letters within the figure indicate significant differences. A significance level of p < 0.05 was chosen.

In line with our findings, previous literature also confirms that certain PNSB strains from the species *Rb. capsulatus, Rps. palustris* and *Rb. sphaeroides* have antimicrobial properties (Kaspari & Klemme, 1977). Chumpol et al. (2017) have performed a similar screening of PNSB on *V. harveyi* and *V. vulnificus.* They isolated 185 PNSB species from shrimp ponds and discovered 10 strains with anti-*Vibrio* potential belonging to the species *Rb. sphaeroides* and *A. marina.* Our research is the first to explore the effect of metabolism and freeze-dried PNSB. From the results of the 96-Well experiment with the freeze-dried PNSB (Figure 1) and the agar spot assay (Figure 2), it can be concluded that PNSB can produce antimicrobial compounds irrelevant of their growth metabolism. To date, there is still a lack of knowledge concerning the nature of the antimicrobial compounds produced by PNSB. It is not known if the compound is species specific or even metabolism specific. Recently, Chumpol et al. (2019) did a preliminary attempt to unravel the structure of a bacteriocin produced by *Rb. sphaeroides.* The study suggested that the compound is a cationic molecule containing NH_2_ groups.

#### 3.2.2 Post-feeding ammonia stress and *Vibrio* challenge test

Additional challenge tests were performed after the feeding trial to validate if dietary inclusion of PNSB could improve the tolerance of shrimp against ammonia and *Vibrio* stress.

Shrimp were exposed to an ammonia stress challenge to explore whether substitution of PNSB in the feed improves the ability of shrimp to withstand physical stress. Research indicates that high ammonia concentrations induce oxidative stress and cause apoptosis in the hepatopancreas of shrimp (Liang et al., 2016). The results for the ammonia stress challenge were promising (Figure 3). Shrimp fed with *Rps. palustris* at both protein substitution levels observed significant higher survival after 48 h (63-75%) compared to the control feed (8%). *Rps. palustris* is known to stimulate the immune parameters when administered live to shrimp ponds (Wang & Gu, 2010). According to Chen et al. (2012), immune parameters decrease after ammonia stress, yet these parameters recover faster when they were initially stimulated by a probiotic. Faster recovery of immune parameters might have contributed to the increased survival after ammonia stress for the *Rps. palustris* fed shrimp. All other diets observed the same survival after 48 h, similar to the commercial feed (*p* > 0.05). Thus, it can be concluded that *Rps. palustris* inclusion in feed enhances the resistance of shrimp against ammonia stress. No effects were observed for *Rb. capsulatus* comparable to the commercial feed.

**Figure 3.**
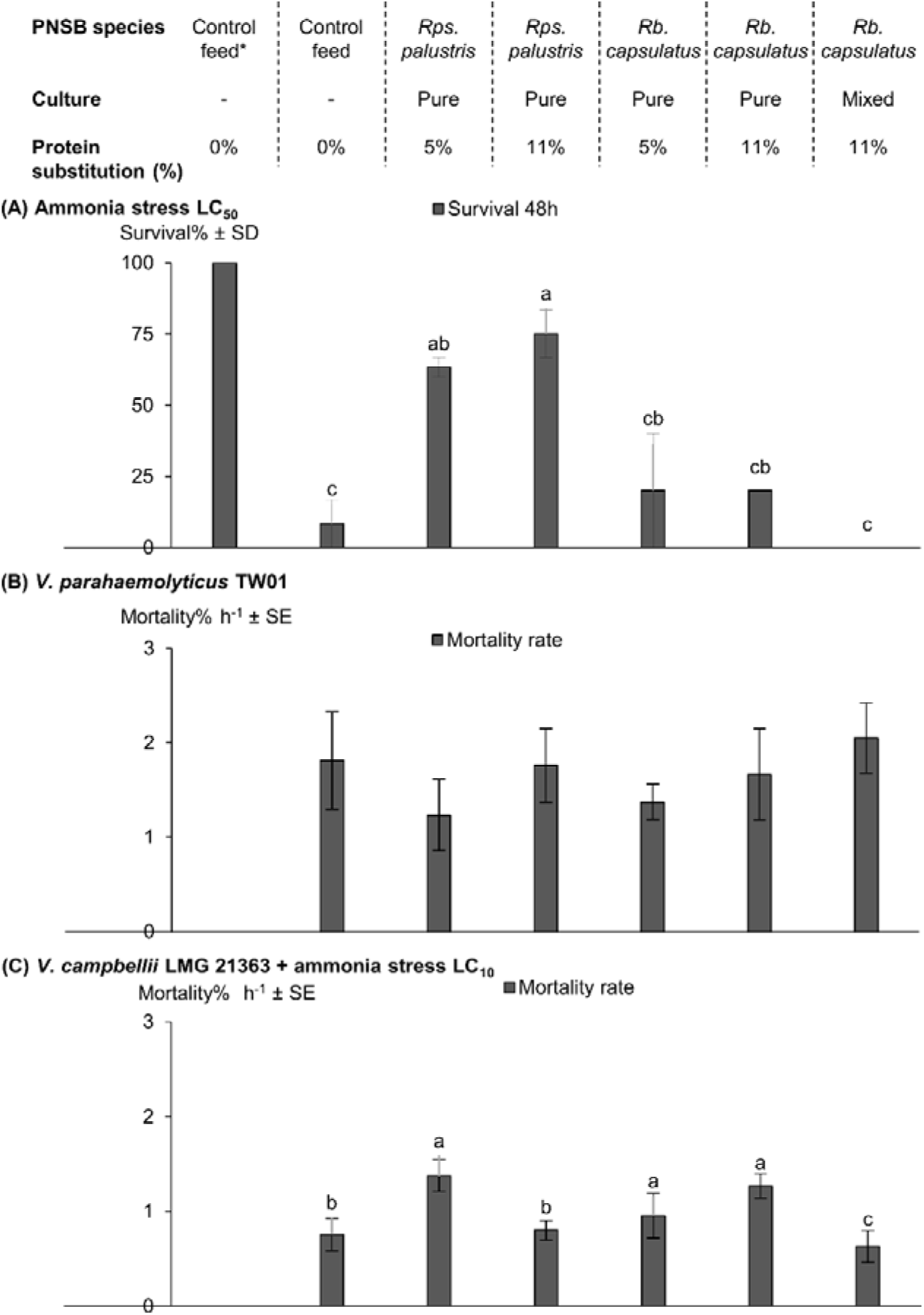
Shrimp survival for (A) the ammonia stress challenge and mortality rate for (B) *Vibrio parahaemolyticus* TW01 challenge and (C) *V. campbellii* LMG 21363 challenge after a feeding trial with a control feed and purple non-sulfur bacteria at two protein substitution levels. *Negative control was shrimp fed with the control feed not exposed to ammonia stress or pathogens. Error bars show standard deviation (SD) or standard error (SE; n=2 for ammonia stress and n=3 for *Vibrio* challenge tests). Letters within the figure indicate significant differences. A significance level of p < 0.05 was chosen.

The shrimp challenged with *V. parahaemolyticus* TW01 (Figure 3.B) were severely affected by the pathogens with a survival of roughly 62% after 14 h. After 39 h, survival was even lower (37%). In terms of mortality rate, there was no significant difference between different diets (Figure 3B). For *V. campbellii* LMG 21363 no mortality was observed before 62 h. For that reason, an extra dose of the pathogen (10^7^ cells mL^-1^) was added to the shrimp to increase the mortality. After 120 h, the survival of all treatments drastically decreased to values between 0-40%. The mortality rate was significantly lower (*p* < 0.05) for *V. campbellii* LMG 21363 than *V. parahaemolyticus* TW01. The lowest mortality rate was observed for the mixed culture of *Rb. capsulatus* followed by the control feed and *Rps. palustris* at a protein substitution of 11%.

## 4 Assessment of PNSB as nutritional feed ingredient

The feeding trial demonstrated that shrimp fed with PNSB contained in the feed had a similar or even an enhanced growth performance relative to a commercial feed (Table 3). Ammonia tolerance of shrimp was even enhanced for shrimp fed with *Rps. palustris*, thereby, reflecting the added-value potential of PNSB. Results of the *in vitro* test showed that PNSB have potential to inhibit *Vibrio* pathogens (Figure 1, Figure 2), yet there was no improved survival against *Vibrio* pathogens for shrimp fed with PNSB in the *in vivo* challenge test (Figure 3). Therefore, it might be possible that PNSB can only suppress the growth of *Vibrio* species if there is a direct PNSB to pathogen interaction as for the *in vitro* tests (Figure 1, Figure 2). Digestion of PNSB will improve the growth performance of shrimp, yet it may not protect shrimp against *Vibrio* pathogens.

Further research is still needed to demonstrate that PNSB can be used as a protein ingredient and protect shrimp against *Vibrio* pathogens. Chumpol et al. (2017) have studied live PNSB administration to shrimp ponds. This improved the shrimp survival against *V. parahaemolyticus.* However, live administered PNSB does not serve as a protein source for shrimp. A potential better application might be the administration of dried PNSB in pure pellets separately from the feed in order to benefit from their full health potential. As such, they will be used as a protein source by shrimp and will also be able to directly interact with *Vibrio-*pathogens. This should of course still be validated along with the dose of PNSB and the frequency of feeding.

In a previous study, we have determined the cost of PNSB produced on wastewater in an open raceway reactor (Alloul et al., 2020). The preliminary cost assessment resulted in a production cost of €1.5 kg dry weight or €3 kg^-1^ protein. As such, the production cost is currently higher than the price of fishmeal (€2 kg^-1^ protein; IndexMundi, 2019)). Prices of probiotics fluctuate around €15-80 kg^-1^. Thus, production costs for PNSB are on par with industrial prices for probiotic. However, if PNSB will be used as added-value protein ingredient, production costs still need to be lower.

## 5 Conclusions

(v) Dietary inclusion of PNSB in shrimp feed resulted in similar and even improved growth performance of shrimp.
(vi) Both freeze-dried and live administration of PNSB suppressed the growth of *V. parahaemolyticus* TW01 and *V. campbellii* LMG 21363 regardless of their metabolism.
(vii) Shrimp fed with *Rps. palustris* are more resistant to ammonia stress.
(viii) PNSB contained in feed enhanced shrimp growth and have the ability to inhibit *Vibrio* pathogens. More research and potentially an alternative way of administering PNSB is needed to benefit from their full health potential.

## Acknowledgements

The authors kindly acknowledge (i) the Research Foundation Flanders (FWO-Vlaanderen) for supporting A.A. with a doctoral fellowship (strategic basic research; 1S23018N), (ii) the Rosa Blanckaert Foundation for supporting A.A with a research grant, (iii) the Belgian Science Policy Office for their support to MELiSSA (CCN5 to C4000109802/13/NL/CP), ESA’s life support system R&D program, which scientifically and logistically supported this study (http://www.esa.int/Our_Activities/Space_Engineering_Technology/Melissa). (iv) Brigitte Van Moffaert and Jorg Desmyter for support during the feeding trials, (v) Koen Blanchaert and Joris Jeursen from VDS (Deerlijk, Belgium) for the feed formulation and the supply of the raw materials. (vi) Dr. Felice Mastroleo from SCK•CEN (Mol, Belgium) for providing *Rhodospirillum rubrum* S 1H (vii) Matthijs Juchem for his assistance in preparing the purple bacteria for the feeding trial, (viii) Gustavo Gomes De Sousa for providing the consortium of aerobic heterotrophic bacteria, (ix) Sarah Lebeer and Camille Allonsius from the Department of Bioscience Engineering, UAntwerp (Antwerpen,Belgium) for their assistance with the agar spot assays.

