## Supplementary Information for "Purple bacteria as added-value protein ingredient in shrimp feed: *Litopenaeus vannamei* growth performance, and tolerance against *Vibrio* and ammonia stress"

### S1. **Synthetic brewery wastewater**

**Table 1** Medium composition for synthetic brewery wastewater

| **Ingredient** | **Mass (mg L^-1^)** |
| --- | --- |
| Beer* | 94 |
| Yeast | 548 |
| Malt extract | 1565 |
| Peptone | 671 |
| (NH4)2SO4 | 1570 |
| MgSO4 | 90 |
| CaCl2 | 75 |
| FeSO4.7H2O | 34 |
| Acetic-C2* | 175 |
| Propionic-C3* | 154 |
| Butyric-C4* | 26 |

*As volume in mL

### S2. **Dose response curve of the ammonia stress challenge**

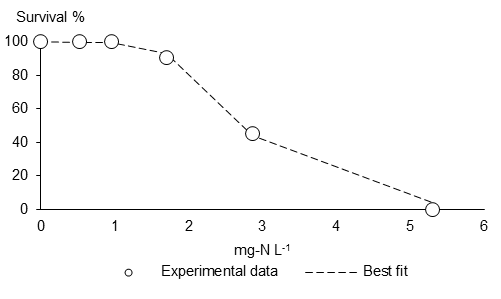

**Figure 1** Dose response curve representing shrimp survival after 48 h of ammonia.

### S3. Growth of PNSB in Marine Broth

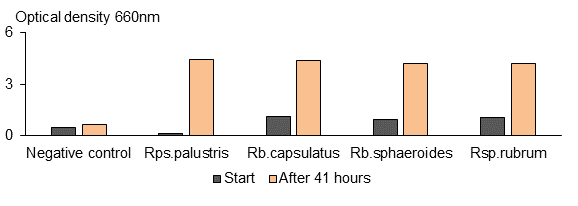

**Figure 2** Chemoheterotrophic growth of *Rhodopseudomonas palustris, Rhodobacter capsulatus, Rhodobacter sphaeroides* and *Rhodospirillum rubrum.*
